# Transcutaneous auricular vagus nerve stimulation increases eye-gaze on salient facial features and oxytocin release

**DOI:** 10.1101/2021.09.12.459983

**Authors:** Siyu Zhu, Yanan Qing, Yingying Zhang, Xiaolu Zhang, Fangyuan Ding, Rong Zhang, Shuxia Yao, Keith Kendrick, Weihua Zhao

## Abstract

**Background:** Transcutaneous auricular vagus nerve stimulation (taVNS) is a non-invasive neuromodulation technique with promising therapeutic potential in the context of epilepsy, pain, and depression and which may also have beneficial effects on social cognition. However, the underlying mechanisms of taVNS are unclear and evidence regarding its role in social cognition improvement is limited.

**Objective:** In order to investigate the impact of taVNS on social cognition we have studied its effects on gaze towards emotional faces using an eye-tracking task and also on release of the neuropeptide oxytocin which plays a key role in influencing social cognition and motivation.

**Methods:** A total of fifty-four subjects were enrolled in a sham-controlled, participant-blind crossover experiment, consisting of two treatment sessions, separated by one week. In one session participants received 30-min taVNS (tragus), and in the other, they received 30-min sham (earlobe) stimulation with the treatment order counterbalanced across participants. Gaze duration towards the faces and facial features (eyes, nose, and mouth) were measured together with resting pupil size. Additionally, saliva samples were taken for the measurement of oxytocin concentrations by enzyme-linked immunoassay.

**Results:** Saliva oxytocin concentrations increased significantly after taVNS compared to sham stimulation, while resting pupil size did not. In addition, taVNS increased fixation time on the nose region irrespective of face emotion, and this was positively correlated with increased saliva oxytocin concentrations.

**Conclusion:** Our findings suggest that taVNS biases visual attention towards socially salient facial features across different emotions and this is associated with its effects on increasing endogenous oxytocin release.

## Introduction

Vagus nerve stimulation (VNS) has been widely used in many clinical conditions for several decades, ranging from neurological disorders to splanchnic diseases, due to the important role of the vagus nerve in communicating with the brain and visceral organs [1,2]. Initially VNS primarily involved a stimulation device invasively attached to the cervical branch of the vagus nerve, and resulting in inevitable risks from the surgery required to implant the device [3]. Recently, transcutaneous auricular vagus nerve stimulation (taVNS) has been developed as a non-invasive alternative with Ventureyra (2000) proposing it as a new therapy for epilepsy [4]. Subsequently, taVNS has been increasingly used to help alleviate symptoms not only of epilepsy but also depression, headache and heart failure [5,6].

Studies in healthy populations have also demonstrated that taVNS can enhance cognitive performance [7], notably in the context of social cognition and emotion recognition [7–9], and enhances attention to direct eye gaze [10]. Thus, taVNS may help to increase attention to salient social cues from faces to aid recognition of both identity and emotion. Importantly, taVNS also has some promising therapeutic effects on disorders with impaired social cognition, including anxiety [11,12], depression [13,14] and post-traumatic stress disorder (PTSD) [15,16]. However, although previous studies suggest that taVNS may improve emotion recognition and attention towards faces, none have directly investigated whether it influences the pattern of gaze directed towards different face emotions using an eye-tracking approach. The main objective of the current study was therefore to investigate whether taVNS influenced attention towards specific salient parts of emotional faces (i.e. eyes, nose and mouth).

Although both preclinical and clinical studies have indicated promising benefits from taVNS, the underlying mechanism(s) involved remain unclear [17]. It has been established that stimulation of the vagus nerve can directly modulate the activation of the brainstem locus coeruleus-norepinephrine (LC-NE) network [18–21]. Increased cortical and hippocampal concentrations of norepinephrine (NE) have been found in rat models via activation of afferent vagal nerve using invasive VNS (iVNS) [19] and iVNS increases firing rates of NE neurons in the locus coeruleus (LC) [22]. Increased pupil dilation has been proposed to be an indirect indication of increased LC-NE activity [17,23,24], although findings on effects of taVNS on pupil dilation in humans have been inconsistent [25–29].

Brainstem NE networks (i.e. LC and nucleus of the solitary tract, NTS) are the primary recipients of vagal nerve afferent fibers [24] and in turn project to limbic and hypothalamic regions controlling emotion and motivation as well as a range of endocrine functions. Notably, two key hypothalamic regions receiving these projections are the paraventricular (PVN) and supraoptic nuclei containing neurons which synthesize the neuropeptide oxytocin (OXT) and regulate its release both into the blood via the posterior pituitary and also within the brain [30,31]. The ascending vagal pathway is therefore likely to influence the release of endogenous oxytocin and indeed evidence from rats models suggests that stimulation of vagus nerve significantly increases short term neuronal activation of both the NTS and PVN [32] and, more directly, plasma oxytocin concentrations have been found to increase immediately after iVNS, even under anesthesia [33]. In addition, both iVNS and taVNS have been demonstrated to increase the activation of the hypothalamus as well as other relevant regions such as the NTS, amygdala, hippocampus and orbital frontal cortex [34,35]. Importantly in the context of observed taVNS effects on social cognition, oxytocin plays a key role in this respect via increasing the salience of social cues [36], and eye-tracking tasks have revealed that oxytocin can bias visual attention towards social stimuli, such as static and dynamic social images [37], as well as emotional faces [38]. Oxytocin is also increasingly being proposed as a potential pharmacotherapy for a variety of psychiatric conditions involving social dysfunction [39]. Based on this evidence, we therefore hypothesized that taVNS may influence social cognition via modulating oxytocin release and we have therefore investigated this by taking saliva samples before and after real or sham stimulation. Both basal and stimulated changes in peripheral OXT can be measured reliably in saliva samples [40].

We hypothesized that any effects of taVNS in altering gaze towards salient facial features during recognition of face emotions may be associated with its modulation of endogenous oxytocin release. We additionally measured pupil dilation, given some evidence that it may represent an index of the effectiveness of taVNS, and hypothesized that any effects of taVNS on pupil dilation would be associated with its behavioral and endocrine effects.

## Material and methods

### Participants

A total of fifty-four healthy adult Chinese university students were enrolled in our study. In an initial interview, all participants reported being free from medical or psychiatric disorders, current or regular medication and did not consume any alcohol, caffeine or nicotine on the day of the experiment. All subjects had normal or corrected to normal vision. Three subjects were excluded due spending insufficient time viewing the face stimuli in the face emotion recognition task (< 2s) and two due to technical problems, leaving 49 participants (32 males, 19.88±1.62 years old) who were included in the final analysis. The study procedures were approved by the ethical committee of the University of Electronic Science and Technology of China and were in accordance with the latest revision of the declaration of Helsinki. The study was also pre-registered as a clinical trial (ClinicalTrials.gov ID: NCT04890457). All subjects provided written informed consent and were financially compensated for their participation.

### Procedure

We conducted a sham-controlled, participant-blind, crossover eye-tracking experiment consisting of two treatment sessions, separated by one week (see Fig. 1 for the protocol). In one session, participants received taVNS (tragus), and in the other they received sham (earlobe) stimulation with the order of treatment counterbalanced across participants. Upon first arrival, each participant completed a number of questionnaires measuring personality traits, including Chinese versions of State-Trait Anxiety Inventory (STAI; [41]), Beck’s Depression Inventory II [42,43], Autism Spectrum Quotient [44], Social Responsiveness Scale [45] and Toronto Alexithymia Scale [46,47]. Participants additionally completed the Positive and Negative Affect Schedule (PANAS; [48]) as a measure of current mood both before and immediately after the experiment in both treatment sessions.

**Fig. 1.**
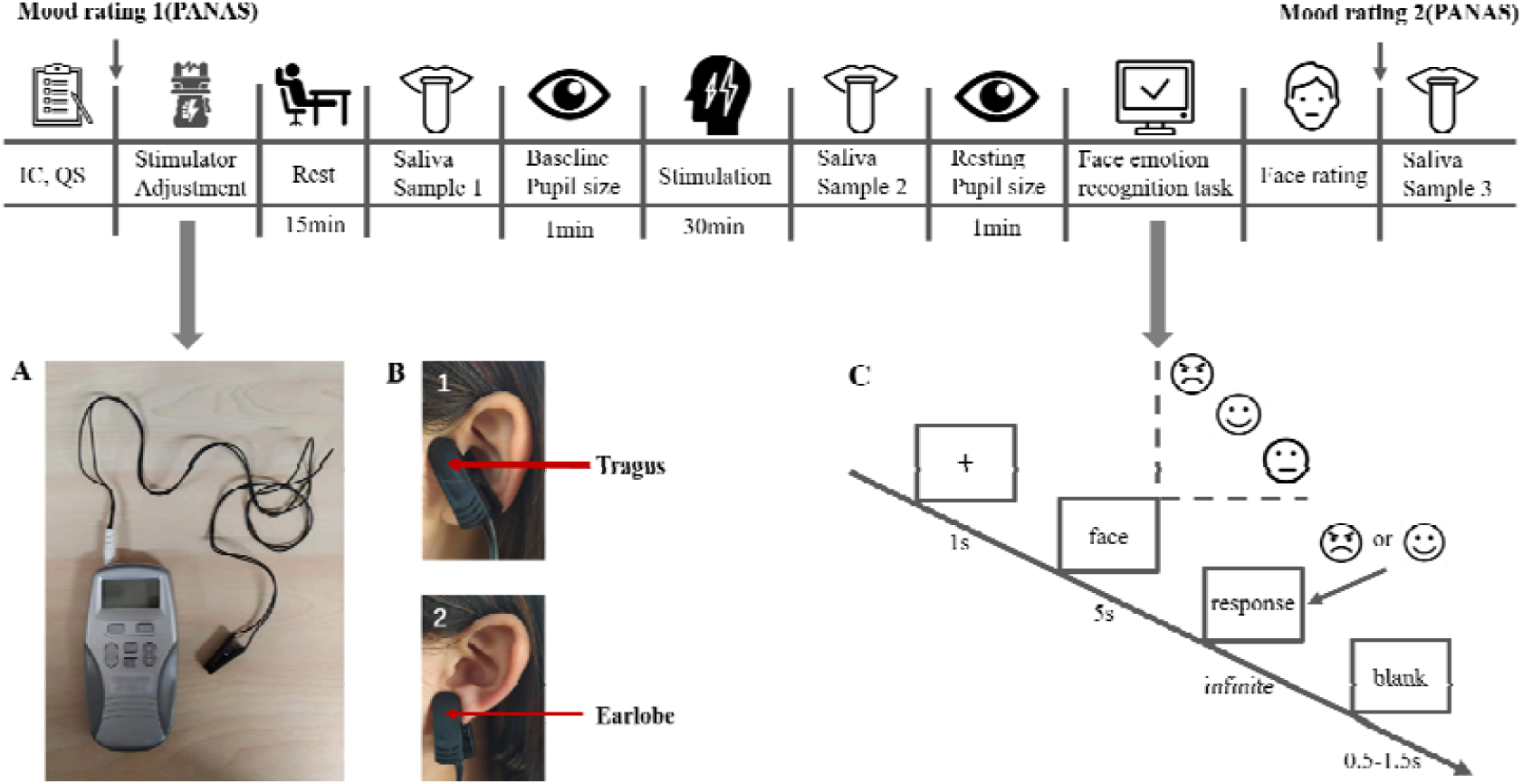
Illustration of procedure. **A**, Transcutaneous electrical stimulator. **B**, Stimulation electrode placement in the left ear: **(1)** location of the taVNS on the tragus; **(2)** location of the sham stimulation on the earlobe. **C**. Trial diagram of face emotion recognition task.

After the completion of questionnaires, participants were asked to sit looking at a display screen in a dimly-lit room to acquire the eye-gaze data (rest and emotion recognition task) using an EyeLink 1000 Plus system (SR Research, Ottawa, Canada). The system was used in monocular mode (right eye) at a sampling rate of 2000 Hz. Participants were instructed to place their head on a chin rest (57 cm from the screen) both for resting pupil size and eye gaze data collection during the face emotion recognition task in order to avoid artifacts due to head movement. A 9-point calibration was performed before each eye tracking procedure to ensure every participant’s pupil was captured (drift correction < 1 of visual angle).

Prior to the formal procedure, participants’ sensitivity to different current intensities of active taVNS or sham stimulation was determined according to their subjective report of whether it induced a “tingling” but not “painful” feeling in line with previous studies [49,50]. The sensitivity to stimulation reported from subjects was different in the real and sham conditions (taVNS: 0.863±0.047 mA; sham: 1.494±0.076 mA) but we considered it more important to match the sensation reported in the two conditions rather than simply match the stimulation current applied. Participants were then asked to rest for 15 minutes in order to avoid any confounding effects of the current stimulation intensity adjustment. Saliva samples were taken immediately after the rest period for measuring baseline oxytocin concentrations (T1), and then 1 minute of resting pupil size was recorded while subjects were instructed to fixate on a white cross centered on the black screen. Subsequently, all participants were asked to do nothing but rest with eyes open during 30 minutes of taVNS or sham stimulation. A second saliva sample (T2) was taken immediately after stimulation and then a second resting pupil size measurement (1 minute) was performed. This was followed by a face emotion recognition task (about 5 minutes). After the task participants were taken to another room and shown the face emotion pictures again and instructed to rate the face emotions in terms of valence (1, very unpleasant, to 9, very pleasant), intensity (1, not strong, to 9, very strong) and arousal (1, not arousing, to 9, very arousing) respectively. When participants had completed their rating a third saliva sample (T3) was collected.

### Transcutaneous auricular vagus nerve stimulation

Electrical pulses (width, 250 μ s) were delivered using a customized transcutaneous electrical stimulator (Fig. 1A, Wuxi Shenping Xintai Medical Technology Co., Ltd) at a frequency of 25Hz with a pre-determined participant-dependent current intensity. On and off periods of stimulation alternated every 30 s, consistent with a number of previous studies [7–9]. In the taVNS condition, an ear-clip electrode was attached to the left tragus (Fig. 1B), which is partly innervated by the auricular branch of vagus nerve [51,52] and its effectiveness in activating vagus nerve projections in the central nervous system has been demonstrated in previous studies [34,49].In the sham condition, the electrode was attached to the left earlobe (Fig. 1B), which was not expected to induce vagus-related brainstem or cortical activation [6].

### Face emotion-recognition eye-tracking task

The present study adopted a face emotion-recognition task (see Fig. 1C). Each trial started with a white fixation cross (1s), which was positioned equivalent to the nasion of the face and on a black background. Face pictures with different emotions (angry, happy and neutral) were randomly presented for 5s in the center of a 17-inch monitor at a resolution of 1024×768 pixels (60Hz) using E-prime 2.0 (Psychology Software Tools, Inc). Subsequently, participant was instructed to indicate whether the face emotion was “angry” or “happy” as soon as possible by pressing two buttons (“F” or “J”) and there was no time limit for responses on each presentation. Following a 500-1000 ms interval, a new trial started again. All face pictures were grayscale images, equalized in size and cumulative brightness. This task included 36 pictures (12 for each emotion: angry, happy and neutral) of 18 Chinese female and 18 Chinese male faces [53]. Two arousal (*t*_(22)_ = 0.426, *p* = 0.673) and intensity (*t*_(22)_ = −0.702, *p* = 0.487) matched face datasets were evaluated by an independent sample for treatment sessions. Accuracy/bias for emotion recognition and response times (RTs), as well as post rating scores (valence, arousal and intensity) were collected.

### Saliva sampling and oxytocin measurement

Saliva samples was collected three times (T1, T2, T3) using Salivette tubes with cotton swab (SARSTEDT, no.51.1534) which were immediately cooled and centrifuged at 1000g for 2 minutes at 4 °C within an hour of collection. Centrifuged saliva was immediately aliquoted into chilled tubes and stored at -80 °C for further oxytocin analysis. All samples were analyzed within 3 months after collection and oxytocin concentrations in 0.5 ml saliva samples measured in duplicate using a commercial enzyme-linked immunosorbent assay (ENZO Oxytocin ELISA kit, Catalog #: ADI-901-153A, Enzo Life Science) and microplate reader (infinite 200 PRO, TECAN Life Sciences). Assay procedures were performed in accordance with the manufacturer’s instructions, including extraction, standards and spiked controls. The extraction step incorporated a 2-fold concentration of saliva samples using a vacuum concentrator (Concentrator plus, Eppendorf, Germany) resulting in a detection sensitivity of 3pg/ml. All samples had detectable concentrations and intra- and inter-assay coefficients of variation were < 9%.

### Eye-tracking and pupillometry recording and data processing

Raw gaze data was initially processed using the Eyelink DataViewer 4.1 (SR Research, Mississauga, Ontario, Canada). For pupil size data, low-pass filtering was performed to remove jittering and linear interpolation was applied to artifacts such as blinks and missing data points when sections of missing data points did not exceed 250ms. To avoid the acute pupil size fluctuation induced by fixating on the screen, we excluded data from the first 5s of the resting pupil size analysis. For the eye gaze data collected during the face emotion recognition task, all raw data with at least 80% gaze weight were analyzed. Fixation duration on each salient facial feature (i.e., eyes, nose and mouth) was calculated using four non overlapping areas of interest (AOI, see Fig. S1, left eye = 3645 pixels, right eye = 3645 pixels, nose = 6955 pixels, mouth = 7420 pixels).

### Statistical analysis

All statistical analyses were performed using SPSS 25.0 (SPSS Inc., Chicago, IL, USA). For behavioral data, response time, response bias (the proportion of recognizing neutral faces as angry or happy), response accuracy (indexed by A prime, for details see Supplementary materials) and post behavioral ratings (intensity, arousal and valence) were analyzed. For response time, two-way repeated ANOVA was performed with two within-subject factors: treatment (taVNS vs. sham) and perceived face emotions (angry vs. happy). For response accuracy and bias, paired t tests were used with treatment (taVNS vs. sham) as a within-subject factor. For post-task ratings, two-way repeated ANOVA was conducted with two within-subject factors: treatment (taVNS vs. sham) and face emotions (angry vs. happy vs. neutral). For saliva oxytocin concentration and resting pupil size data, two-way repeated ANOVAs were performed with treatment and time point [saliva OXT: baseline (T1), after 30 minutes stimulation (T2), end of the experiment (T3); resting pupil size: baseline, after stimulation] as two within-subject factors.

For eye-tracking data, percentage of total fixation duration in the face emotion-recognition task was calculated by dividing fixation time spent viewing each AOI (eyes, nose and mouth) by the time spent viewing the whole face and a three-way repeated ANOVA was conducted with three within-subject factors: treatment, face emotion (angry, happy and neutral) and AOIs (eyes, nose and mouth). Bonferroni correction was applied to all post-hoc tests. In addition, Spearman correlation analysis was performed to investigate the relationship between the percentage of increased oxytocin concentrations (after stimulation/baseline × 100) and the percentage of increased fixation time on facial features (percentage of fixation time after taVNS/ percentage of fixation time after sham stimulation ×100).

## Results

There were no significant differences on participants’ positive and negative mood scores (PANAS) were found before and after tasks for taVNS and sham treatment (see Table 1). No significant effects of taVNS were observed on response times, accuracy, bias or post-task ratings of face emotions (all *p*s ≥ 0.132, for more details see in Supplementary Results).

**Table 1.** Positive and Negative Affect Schedule (PANAS) scores in before and after taVNS and sham treatment (mean ± SEM)

|  | taVNS | Sham | *t*-Value | *p*-Value |
| --- | --- | --- | --- | --- |
| Pre-task |  |  |  |  |
| Positive scores | 25.592 ± 0.906 | 25.102 ± 0.806 | 0.628 | 0.533 |
| Negative scores | 14.714 ± 1.043 | 15.735 ± 1.106 | -1.468 | 0.149 |
| Post-task |  |  |  |  |
| Positive scores | 24.592 ± 0.922 | 24.286 ± 1.008 | 0.398 | 0.692 |
| Negative scores | 14.347 ± 1.083 | 14.429 ± 1.193 | -0.137 | 0.891 |

### Effects of taVNS on oxytocin release

A two-way repeated ANOVA showed that there was a significant treatment × time point interaction effect (*F*_(2, 96)_ = 11.034, *p* < 0.001, partial □^*2*^= 0.187). A Bonferroni-corrected post-hoc analysis demonstrated that taVNS significantly increased saliva oxytocin concentrations immediately after 30 minutes of stimulation compared to baseline (*p* < 0.001, Cohen’s *d* = 0.745) while sham stimulation did not (*p* = 0.638). There was no difference between baseline oxytocin concentrations in the taVNS and sham stimulation conditions (*p* = 0.270, see Fig. 2).

**Fig. 2.**
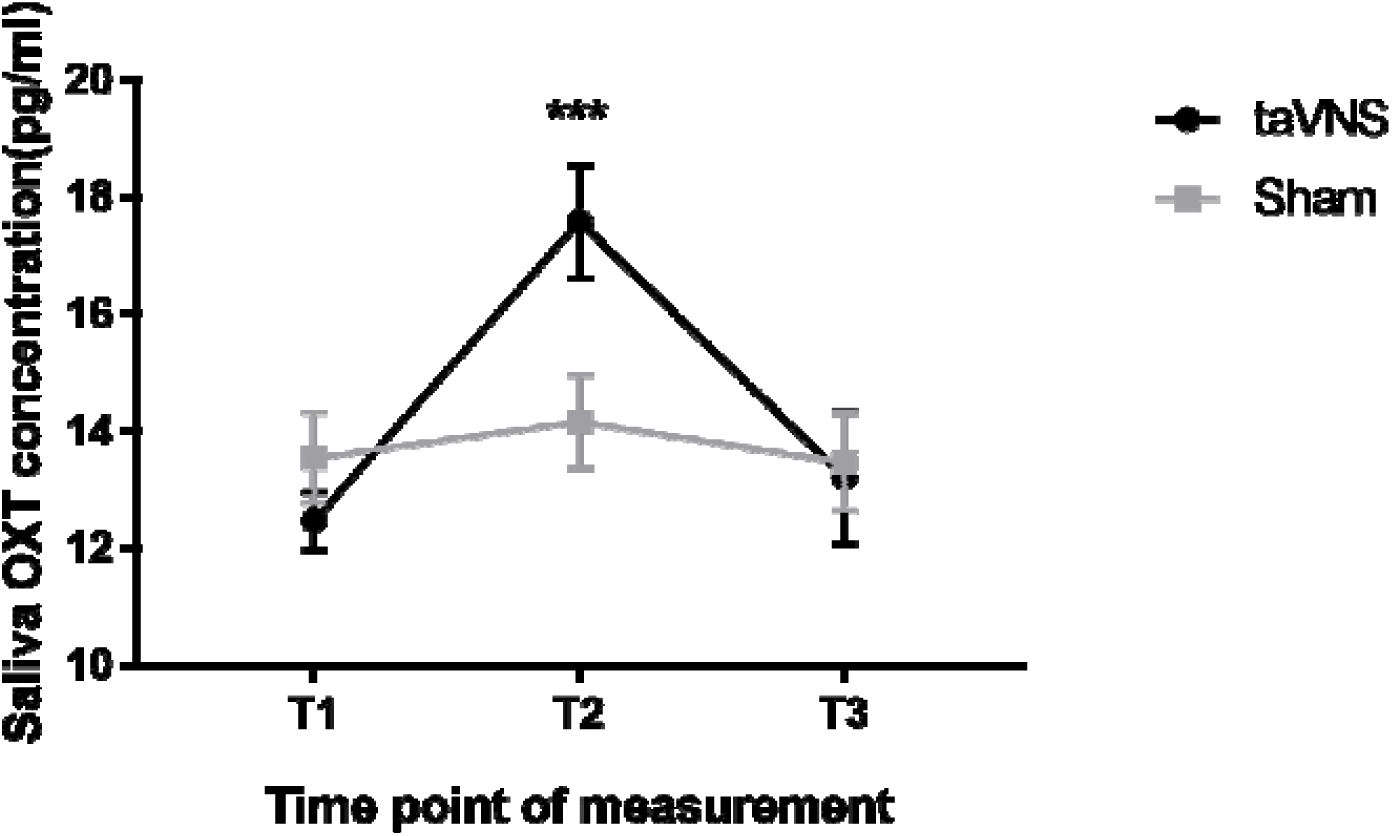
Saliva oxytocin concentrations (mean ± SEM) at three time point under taVNS and sham treatment. T1: baseline measurement; T2: measurement right after 30 minutes taVNS/sham stimulation; T3: measurement at the end of experiment. \*\*\**p* < 0.001.

### Effects of taVNS on percentage of fixation duration on face regions

To investigate whether taVNS would influence the percentage of fixation time on the three salient face regions, a three-way repeated ANOVA with treatment, face emotions and regions as within-subject factors was conducted. Results revealed an interaction effect between treatment and face regions (*F*_(2, 96)_ = 4.115, *p* = 0.028, partial □^*2*^ = 0.079) and post-hoc analysis with Bonferonni correction showed that participants spent proportionately more time looking at the nose region after taVNS compared to the sham stimulation condition and irrespective of face emotions (*t*_(48)_ = 2.143, *p* = 0.037, Cohen’s *d* = 0.220, a heat map of fixation duration for one face emotion is shown in Fig. S2), but not for the eyes (*t*_(48)_ = −1.425, *p* = 0.161), or mouth regions (*t*_(48)_ = −1.650, *p* = 0.106) (Fig. 3A).

**Fig. 3.**
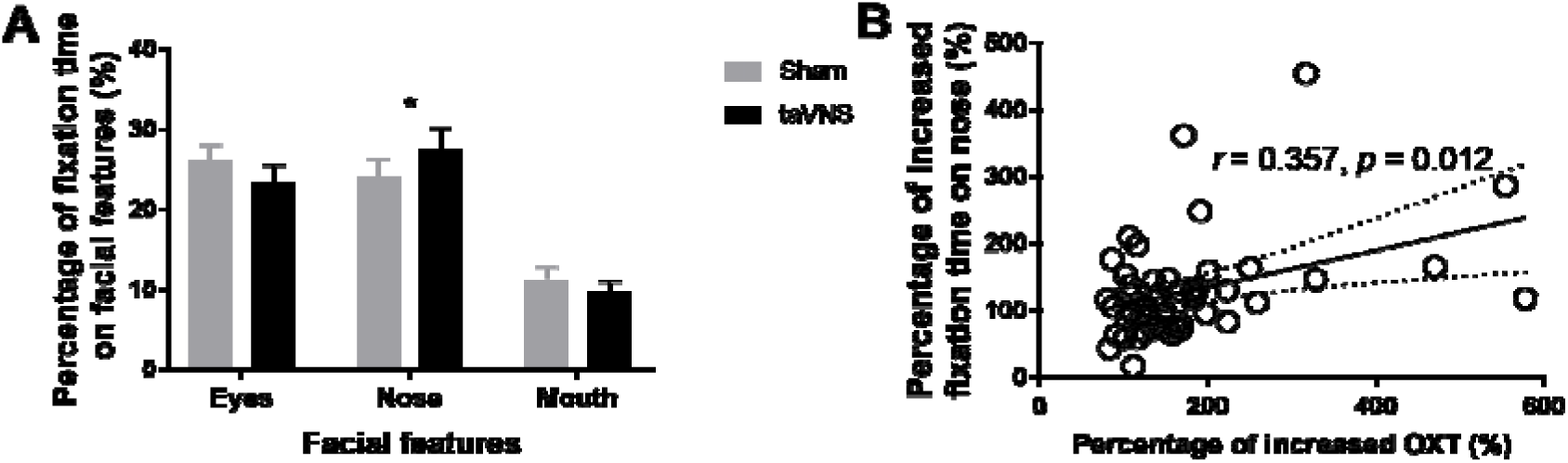
**A**, Percentage of fixation time (mean ± SEM) on different facial features (eyes, nose and mouth) across emotions under taVNS and sham treatment. **B**, Correlation analysis between increased fixation time on nose and increased OXT level after taVNS. \**p* < 0.05.

### Associations between fixation durations and oxytocin concentrations

A significant positive correlation between increased proportion of fixation time on nose and increased oxytocin concentrations after taVNS was found via Spearman correlation analysis (*r* = 0.357, *p* = 0.018, see Fig. 3B), suggesting a positive association between taVNS effects on increased viewing of the nose region and those on increasing oxytocin release.

### Effects of taVNS on pupil size

For analysis of taVNS effects on pupil size, a two-way repeated ANOVA was applied with treatment and time point as within-subject factors. There was no main effect of treatment (*F*_(1, 48)_ = 0.119, *p* = 0.732) or interaction effect with treatment (*F*_(2, 96)_ = 0.332, *p* = 0.567). A marginal main effect of time point was found (*F*_(1, 48)_ = 3.997, *p* = 0.051, partial □^*2*^ = 0.077), indicating a decrease in pupil diameter from baseline measurement to 30 minutes following stimulation in both taVNS and sham conditions (Cohen’s *d* = 0.095).

## Discussion

The present study investigated the impact of taVNS on patterns of eye gaze during face emotion recognition using an eye-tracking task and also on peripheral release of the neuropeptide oxytocin and on pupil diameter. Results revealed that taVNS promoted increased oxytocin release but did not affect resting pupil size. In addition, taVNS increased fixation time spent on the nose region irrespective of face emotion. Furthermore, increased fixation time on the nose region was positively correlated with increased saliva oxytocin concentrations.

We found that taVNS compared to the sham condition increased the proportion of time spent viewing the nose region irrespective of face emotions. Preferential scanning of the nose region during discrimination of own-race faces in Asian subjects has been reported in several previous studies [54–57]. Thus the nose seems to be a particularly social salient feature for Asians to process faces and emotions in social situations and this nose-centric strategy may help facilitate holistic processing of faces [58]. It is unlikely that our finding was influenced by the location of the fixation cross on the screen prior to each face presentation since this was deliberately positioned at the level of the nasion (i.e. above the AOI we used for the nose region). It is likely that this increased viewing time of the nose region may aid recognition of both the identity and emotions being expressed by faces. In the current study, we deliberately used only a small number of easily recognized face emotions to focus primarily on patterns of gaze, and recognition accuracy was very high > 98% and response times very fast. Such high accuracy represents a ceiling effect which precluded any realistic assessment of whether taVNS improved face recognition accuracy as a result of increased fixation of the nose region. Indeed, previous studies demonstrating face emotion recognition improvements following taVNS have primarily found them for difficult stimuli [32].

As predicted, we found that taVNS significantly increased endogenous peripheral oxytocin concentrations measured in saliva. While altered peripheral concentrations of oxytocin may not necessarily reflect altered concentrations within the brain, the association between salivary and cerebroventricular concentrations in humans is reasonable [59]. This is the first evidence for taVNS induced oxytocin release in humans and is consistent with a previous finding that iVNS increased plasma oxytocin concentrations in anesthetized rats [33]. Thus, taVNS may increase the activity of hypothalamic neurons containing oxytocin via an ascending pathway from the auricular branch of vagus nerve and its brainstem target regions projecting to the hypothalamus [34,51]. Notably the hypothalamic paraventricular nucleus (PVN), which contains many oxytocin neurons both receives projections from the vagus nerve (VN) and sends projections to the dorsal vagal complex [31,60]. Importantly, taVNS-evoked oxytocin release was positively associated with increased time spent viewing the nose region which supports this as a possible mechanism whereby vagal stimulation influences social cognition. Further support for this possibility comes from a number of studies reporting that exogenous administration of oxytocin in humans via nasal spray also has potent effects on many aspects of social cognition [39]. Indeed, the magnitude of increased oxytocin concentrations following taVNS are very similar to those found after a single treatment with intranasal oxytocin at 24IU [61]. Furthermore, exogenous oxytocin treatment has been reported to improve social deficits in autism spectrum disorder and a recent study in a mouse model of autism has found that treatment of their social impairment is dependent on an interaction between oxytocin and vagus nerve function [68]. Taken together these findings suggest that taVNS facilitation of endogenous oxytocin release may have therapeutic benefits in the context of disorders with social dysfunction.

While pupil dilation has been considered as a potential biosensor for successful VNS, particularly following findings in animal model studies [62,63], our results show no effect of taVNS on resting pupil size in humans in line with a number of other studies [27–29,64]. There are several reasons why we may have failed to detect pupil dilation changes under taVNS relative to the sham condition: (1) the transcutaneous protocol used was insufficient to stimulate the vagus nerve sufficiently, perhaps due to the stimulation parameters (pulse width, frequency and amplitude) which still need further optimization [6,65]; however our current study did find both eye-gaze and oxytocin concentration changes after the stimulation, which suggest that the taVNS parameters chosen do have functional effects; (2) the procedure used to measure pupil dilation (i.e., the way we defined baseline pupil size and selection of time points) might also have influenced the results. For example, the effects of taVNS on pupil dilation could only be quite transient [25,26]. Indeed, two recent studies have reported that short trains of taVNS induced pupil dilation but which rapidly returned to baseline level [25,26], and so failure to detect pupil size changes after a long period of taVNS stimulation may simply reflect this transience. Thus, it may be useful to collect a resting pupil sizes more frequently after the onset of taVNS in future studies.

## Conclusions

In summary, the present study demonstrates that taVNS as a developing non-invasive technique, could effectively increase fixation time on the nose region of faces across different emotions and increase peripheral oxytocin release. Furthermore, the impact of taVNS on increasing visual attention towards the nose was positively associated with its effects on increasing oxytocin concentrations. These findings suggest that taVNS may be a promising therapeutic treatment for enhancing social cognitive functions and oxytocin release in clinical conditions, especially disorders with social deficits (i.e., autism spectrum disorder, schizophrenia, depression and anxiety).

